# Promoter Choice: Who Should Drive the CAR in T Cells?

**DOI:** 10.1101/2020.04.27.063743

**Authors:** Ali Hosseini Rad SM, Aarati Poudel, Grace Min Yi Tan, Alexander D. McLellan

**Affiliations:** Department of Microbiology and Immunology, University of Otago, Dunedin, New Zealand

**Author notes:** Grace Min Yi Tan and Aarati Poudel contributed equally to this work. To whom correspondence should be addressed: Alexander D. McLellan; Department of Microbiology and Immunology, University of Otago, Dunedin 9010, Otago, New Zealand;.

**Keywords:** long RNA, chimeric antigen receptor (CAR), Mcl-1, Immunotherapy, Promoter

## Abstract

Chimeric antigen receptor (CAR) T cell therapy is an effective treatment for B cell malignancies, with emerging potential for the treatment of other hematologic cancers and solid tumors. The strength of the promoter within the CAR cassette will alter CAR-polypeptide levels on the cell surface of the T cell – impacting on the kinetics of activation, survival and memory cell formation in T cells. In addition to the CAR, promoters can be used to drive other genes of interest to enhance CAR T cell function. Expressing multiple genes from a single RNA transcript can be effectively achieved by linking the genes via a ribosomal skip site. However, promoters may differ in their ability to transcribe longer RNAs, or could interfere with lentiviral production, or transduction frequencies. In this study we compared the ability of the strong well-characterized promoters CMV, EF-1, hPGK and RPBSA to drive functional expression of a single RNA encoding three products: GFP, CAR, plus an additional cell-survival gene, Mcl-1. Although the four promoters produced similarly high lentiviral titres, EF-1 gave the best transduction efficacy of primary T cells. Major differences were found in the ability of the promoters to drive expression of long RNA encoding GFP, CAR and Mcl-1, highlighting promoter choice as an important consideration for gene therapy applications requiring the expression of long and complex mRNA.

## Introduction

Promoters are of critical importance for expressing optimal levels of the transgene in CAR T cells for the production of functional proteins (1–5). It is also clear that high expression of the CAR can result in antigen-independent CAR signaling, resulting in T cell exhaustion and sub-optimal anti-tumor responses, or exacerbate the inappropriate recognition of tumor antigen on self-tissue (1, 2). In addition, controlling CAR T cell signaling is critical for proper memory cell formation (6). Because surface expression of the CAR may be limited by RNA levels, the choice of promoter is critical (1, 2).

There have been limited studies that directly compare the efficiency of different promoters for driving long mRNA comprising multiple genes within CAR T cells (1, 2, 7). Recent studies investigating promoter performance in mouse or human T cells were usually limited to either the CAR, a single gene of interest alone, or single fluorescent reporter genes of limited size (1, 2, 7–9). For the generation of lentiviral particles for transduction, using multiple internal promoters or internal ribosome entry sites (IRES) for multiple genes may interfere with transcription or reverse transcription of RNA genomic, impacting upon lentiviral particle titre, and / or on the efficiency of integration into the target cell (8, 10). Therefore, strategies that employ single promoters to drive multiple genes may be preferred for CAR T cell engineering (9).

Although all current, clinically-approved second and third generation CAR T cells rely on the expression of a single gene encoding a single polypeptide, it may be advantageous to express longer RNA containing the CAR, together with one or more genes of interest. For example, endogenous growth factors or membrane bound or secreted cytokines could improve T cell expansion and survival (6, 11). Alternatively, markers of transduction efficiency or death switches could be incorporated into the CAR element (4, 12–14). Promoter choice for such applications is crucial to obtain optimised gene expression of multiple, linked genes.

Because requirements for driving short versus long RNA might be distinct in CAR T cell genetic elements, we investigated the ability of several promoters to drive an extended downstream genetic sequence comprised of GFP, anti-Her2-CAR and an additional cell survival gene Myeloid leukemia cell differentiation protein (Mcl-1), an anti-apoptotic Bcl2 family member. Mcl-1 aids in T cell development, mitochondrial function and lifespan and appears to a suitable candidate for enhancing CAR T cell performance (15, 16). Mcl-1 inhibits the action of pro-apoptotic BIM / BAK / BAX at the mitochondrial membrane and is expressed throughout T cell differentiation and is essential for memory T cell formation (16–20).

The individual elements were tested at protein level and for functional activity. The results demonstrated clear differences in the ability of these internal promoters to drive expression of multiple CAR-cassette associated transgenes.

## Material and Methods

### Plasmid construction

The third-generation lentiviral vector pCCLsin.cPPT.hPGK.GFP.WPRE (pCCLsin) and VSV-G-based packaging plasmids were a kind gift from Prof. Dr. Naldini and have been described elsewhere (21). The anti-Her-2 CAR FRP5, anti-CD19 CAR FMC63 (with -EQKLISEEDL-c-myc tag between scFv and CD8 hinge) and codon-optimized human Mcl-1 (cop-Mcl-1) were synthesized as gene blocks (IDT Technologies). Sap I type IIs restriction enzyme cloning was utilized for scarless assembly of the eGFP-P2A-CAR-P2A-Mcl-1. This cassette was then cloned into the BamHI and SalI sites of the pCCLsin (Figure 1a). Promoters were amplified with 5’ EcoRV and 3’ BamHI sites from respective plasmids: CMV from pcDNA3.1(-), EF-1 from Sleeping Beauty (pSBbiRP) and RPBSA from Sleeping Beauty (pSBtet-GP) and ligated upstream of the GFP-CAR-mcl1 cassette. Codon optimized Leucine Zipper CD95 (LZ-CD95L) gene was synthesized by IDT with EcoRI and BamHI sites and cloned into pcDNA3.1(-) (Addgene #104349).

**Fig. 1.**
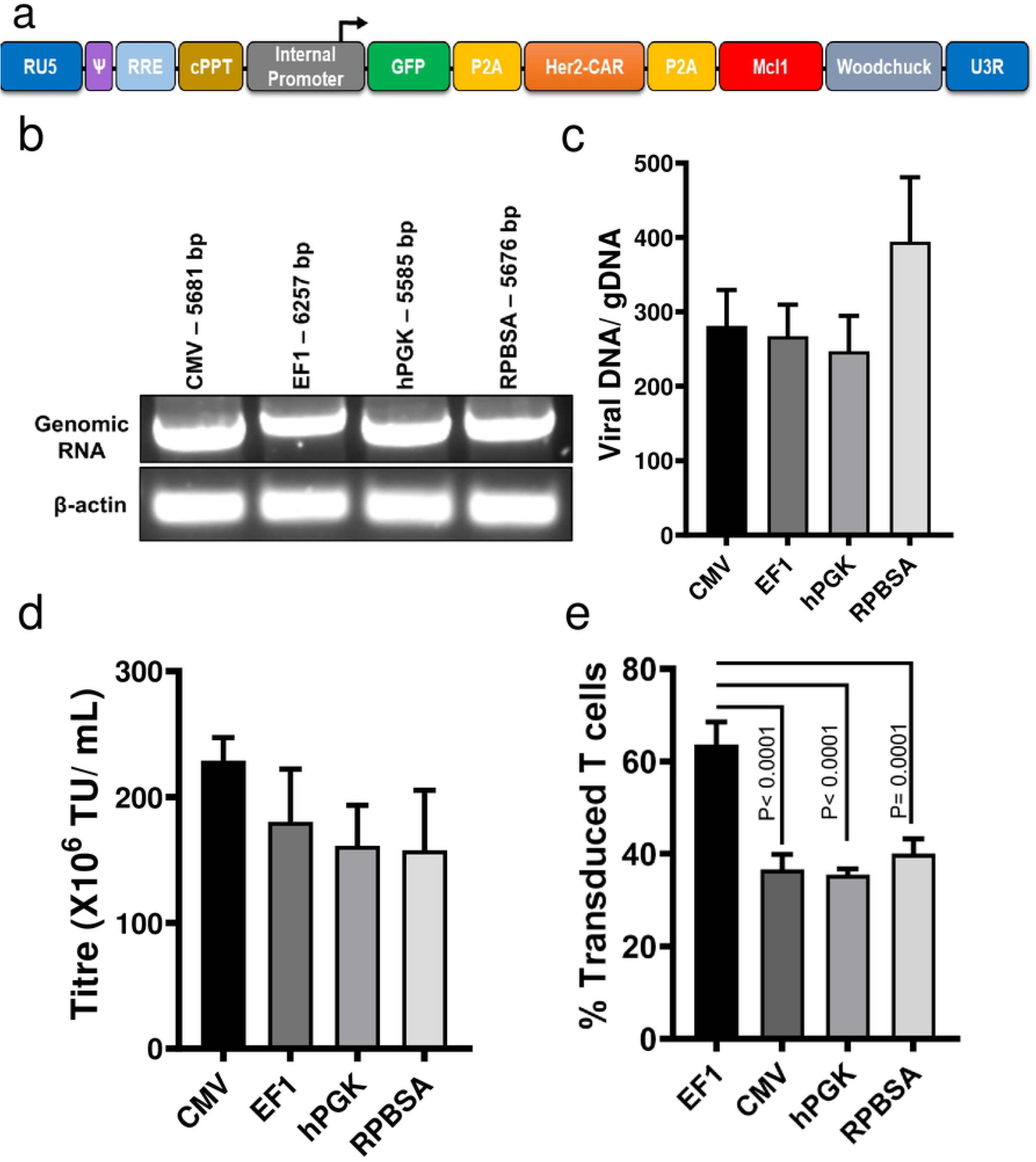
The effect of internal promoters in producing functional lentiviral particles **a)** Schematic illustration of the pCCLsin backbone bearing four different internal promoters (CMV, EF-1, hPGK and RPBSA) for driving a long RNA consist of GFP-P2A-Her2CAR-P2A-Mcl-1 **b)** HEK293T cells were transfected with lentiviral constructs containing different promoters along with packaging plasmids. At 24 h post transfection, total RNA was extracted and 1 μg of RNA was converted to cDNA. PCR was carried out using specific primers binding to PPT and woodchuck region. Agarose gel electrophoresis displays the PCR product band of each construct. Lower band displays the PCR product of β-actin serving as a loading control. **c)** Ratio between integrated viral cassettes to gDNA 48 h post-transduction. Genomic DNA was extracted from cell lysates and qPCR was performed using Gag for integrated lentivirus, β2 microglobulin for gDNA and β-actin as a housekeeping gene. **d)** Comparing the viral titration of four different constructs by analyzing the percentage of GFP expression in HEK293T cells using flow cytometry. Bar graph values represent the titre unit/mL (TU/mL) from three independent repeats **e)** Transduction efficacy of primary T cells for four lentivectors. CD3/ CD28 stimulated human primary T cells were transduced at MOI 40 and cells were analyzed for GFP expression 72 h after transduction by flow cytometry. Dead cells were excluded with Zombie NIR viability dye gating at analysis. Bar graph values represent the mean values ± SD from three different repeats.

### Cell culture

Cell lines were cultured in a humidified atmosphere at 37°C, 5% CO_2_ (or with 8% CO_2_ for LV-Max and Expi293F). Human embryonic kidney 293T (ATCC CRL-1573) and MCF-7 (ATCC HTB-22) cell lines were cultured in high glucose Dulbecco’s Modified Essential medium (DMEM) supplemented with 10% fetal bovine serum (FBS; Pan-Biotech GmbH), penicillin (100 U/mL) and streptomycin (100 μg/mL) (Gibco). MCF-7 and HEK293T cells were transfected using Lipofectamine 3000 according to manufacturer’s protocol.

Human peripheral blood mononuclear cells (PBMC) were isolated from healthy donors with consent (Ethics H18/089). Frozen PBMCs were thawed and then rested overnight in T cell expansion media (Thermofisher #A1048501) supplemented with 50 U/mL of hIL-2 (Peprotech, #200-02), L-glutamine and 10 U/mL penicillin and streptomycin (Gibco). prior to CD4 and CD8 T cells isolation using EasySep Human T cell isolation kit (STEMCELL Technology, #17951). Isolated T cells were activated with Dynabeads Human T-Activator CD3/CD28 (ThermoFisher, # 111.32D).

### Lentiviral production, titration and T cell transduction

Lentviral production and titration was carried out using LV-Max Viral production system (ThermoFisher #A35684) according to manufacturer’s protocol. HEK293T cells were transduced at MOI 2:1 with 8 μg/mL of polybrene (Sigma-Aldrich). One day before transduction, plates were coated with 40 μg/mL retronectin (TAKARA, # T100A/B) overnight at 4°C, blocked with 2 % FBS/PBS for 15 min, before adding LV at 40:1 MOI to the plate; followed by spinoculation at 800 ×g for 2.5 h at room temperature. After 48 h of activation with a 1:1 ratio of CD3/CD28 Dynabeads, T cells were added to virus-coated wells and spun at 500 ×g for 5 min. The next day, T cells were debeaded and cultured in media plus 50 U/ mL of hIL-2. Media was changed with fresh medium supplemented with 50 U/mL hIL-2 every three days.

### RNA extraction, long cDNA synthesis and RT-PCR

Total cellular RNA (containing viral genomic RNA) was extracted 48 h after transfection using NucleoSpin RNA Plus kit (Macherey-Nagel, Germany) according to the manufacturer’s protocol. Then RNA was reverse transcribed using PrimeScript™ RT Reagent Kit (Takara Bio, USA) according to manufacturer’s protocol RT-PCR was performed using internal primers PPT-Fwd: GGGTACAGTGCAGGGGAAAG and Woodchuck-Rev: AAGCAGCGTATCCACATAGCG for comparison with β-actin Fwd: CTTCCTTCCTGGGCATG and β-actin-Rev: GTCTTTGCGGATGTCCAC.

### Quantification of gDNA/ integrated viral DNA ratio

At 48 h post transduction, integrated lentiviral DNA was quantified by extracting genomic DNA using Qiamp DNA Mini kit (Qiagen, Germany) and the ratio of viral genome: human gDNA were estimated using qPCR via Luna Universal qPCR Master Mix (New England Biolabs) using designed primers Gag-Fwd: GGA GCT AGA ACG ATT CGC AGT TA, Gag-Rev: GGT TGT AGC TGT CCC AGT ATT TG TC, PBS-Fwd: TCT CGA CGC AGG ACT CG; PBS-Rev: TAC TGA CGC TCT CGC ACC, and β-actin forward and reverse primers described above. All reactions were run in triplicate and were presented as mean ± SD.

### Western blot

Cell lysates were prepared using RIPA lysis buffer and blotting carried out using mouse monoclonal anti-EGFP antibody (Abcam, #ab184601), rabbit anti-human Mcl-1 (Abcam, #ab28147), biotin anti-c-myc (Biolegend #908805). Mouse monoclonal β-actin primary antibody (Sigma-Aldrich #A2228) was used as loading control. The membrane was scanned using an Odyssey Fc imaging system (Licor, Germany) and analyzed using Image Studio Lite software.

### Mitochondrial Membrane Potential Assay (TMRE)

Transduced T cells were incubated overnight with 1 μg/mL of LZ-CD95L, then 4 μM TMRE (Invitrogen) was added at 37°C for 30 min. GFP positive cells were analyzed by flow cytometry.

### Cytotoxicity and cytokine release assay

Luciferase-based cytotoxicity assay was carried out for Her2 and CD19 CAR T cells as previously described (22) at a 10:1 ratio of effector to target cells of using Firefly Luc One-Step Glow assay (ThermoFisher #16197). For analysis of cytokine release, CAR T cells were added to target cells in a 2:1 ratio. IL-2 and IFN-γ concentration secreted in cell supernatant were measured using sandwich ELISA according to manufacturer’s protocol (BD Biosciences, USA). Plates were read on a Varioskan Lux multimode microplate reader (ThermoFisher, USA).

### Flow cytometry

CAR T cells were stained with biotin anti-c-myc antibody (Biolegend #908805) detected with Streptavidin-Brilliant Violet 421 (Biolegend #405225). Antigen experienced CAR T cells were stained for CD69 expression using APC-conjugated anti-human CD69 antibody (Biolegend #310910). Flow cytometric data was acquired using a BD LSRFortessa with BD FACSDiva software. Data was analysed with FlowJo v10.6.2 software. Cells were subject to FSc and SSc doublet discrimination and dead cells were excluded from analysis using Zombie NIR viability dye (Biolegend #423106).

### Statistical analysis

All experiments were carried out at least three times, presented as mean ± standard deviation (SD) and analyzed by student T test and ANOVA test with Bonferroni post-test correction. The P values of ≤0.05 were considered statistically significant. (* P<0.05, ** P<0.01, *** P<0.001, **** P<0.0001)

## Results

### Compatibility of the promoter systems with a third-generation lentiviral system

The four promoters were chosen based on their widespread use in the literature and documented ability to drive high level expression of transgenes in either lentiviral vectors, or in Sleeping Beauty transposon vectors (1, 8, 9, 21, 23). Each of the four promoters were cloned upstream of the series of P2A-linked genes comprised of GFP, the FRP5 anti-Her2 CAR followed by human Mcl-1 (Fig. 1a), a Bcl2 family member – the latter gene as a strategy to protect CAR T cells against activation-induced cell death (AICD). A first consideration for the choice of internal promoter driving transgenes within lentiviral systems is the effect on viral titration and transduction efficacy. Generally, there is a difference in the degree of transcriptional interference between the internal promoters and the promoter driving expression of genomic RNA, resulting in a lower number of full-length genomic RNAs particularly when the CMV or EF-1 promoter is being used (10, 24). However, similar levels of full-length transcripts were obtained using all constructs, as assessed by RT-PCR carried out with primers binding to cPPT and woodchuck regions (Fig. 1b). The effect of internal promoter interference with provirus production was estimated using qPCR. The ratio between integrated cassette to gDNA did not show significant differences among constructs (P≤0.05, Fig. 1c), suggesting that the selected promoters do not affect reverse transcription or integration steps.

Next, we determined if the choice of internal promoter affects titre and transduction of primary T cells. As shown in Fig. 1d, constructs containing any of the four promoters were able to produce similar viral titres, as determined by transduction of the GFP marker into HEK293T cells. To determine if the sequences of internal promoters affected primary T cells transduction, we transduced primary T cells obtained from different donors and analyzed for GFP expression by flow cytometry three days later. EF-1 gave superior transduction efficacy compared to the other three promoters (P≤ 0.0001, Fig. 1e).

### Promoter comparison for long and complex gene expression

To determine if the promoters differed in their ability to transcribe individual gene products within a long gene, the expression of individual genes were assessed in HEK293T and primary T cells. From the data obtained with HEK293T, CMV and EF-1 were superior to hPGK and RPBSA in producing all three products (Fig. 2). We next examined the strength of the four promoters in primary T cells by analyzing GFP and CAR expression. As it is shown in Fig. 3, EF-1 shows stronger expression of GFP and Her2 CAR than the other promoters. CMV was weaker in primary T cells as compared to HEK293T cells. This could be due to the differences in the transcriptome of both cell types and/or the different techniques that have been used to measure the protein level.

**Fig. 2.**
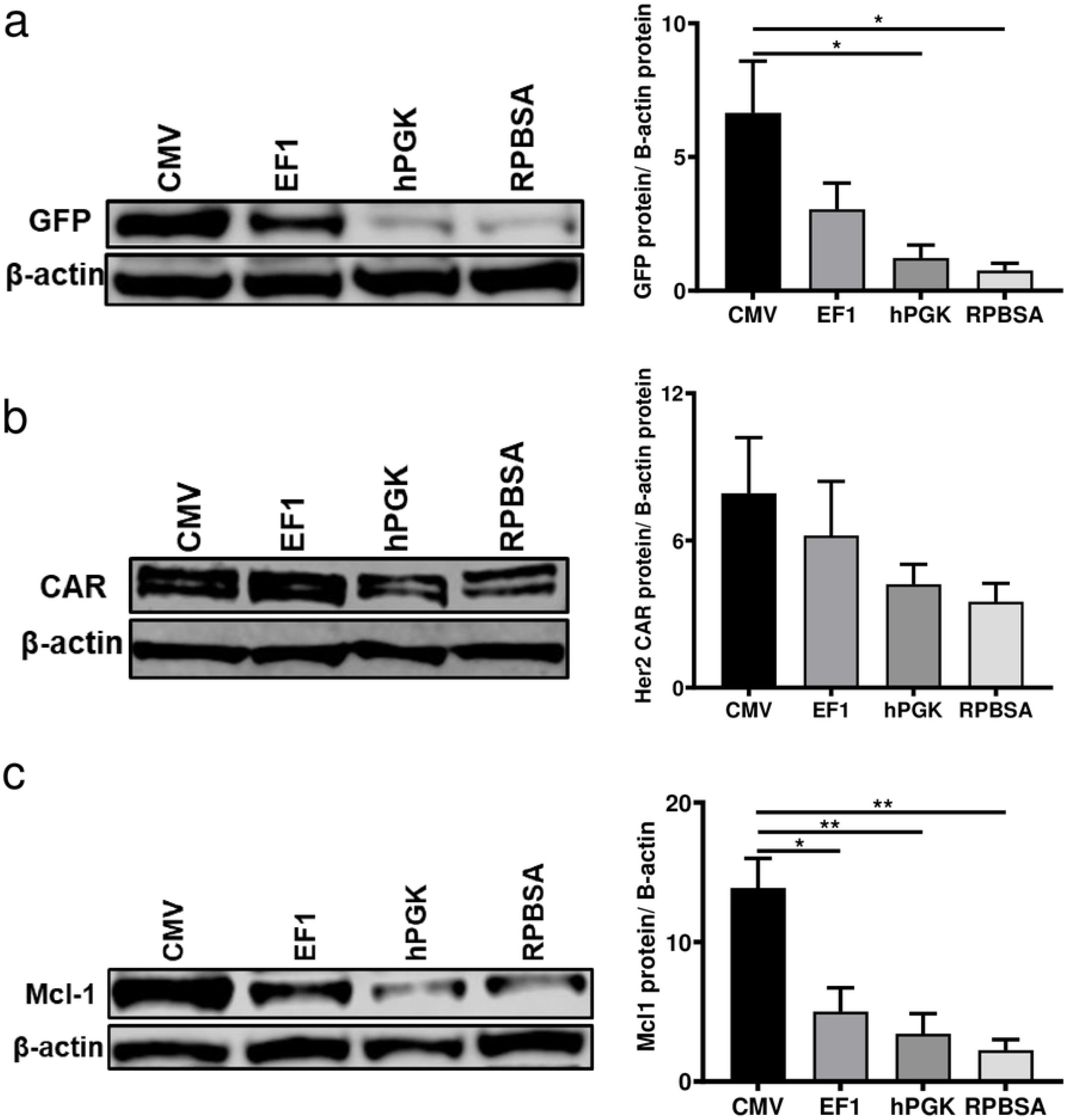
Protein expression from four different constitutive promoters driving long mRNA. Transfected HEK293T cells were lysed with RIPA buffer and processed for immunoblotting using antibodies to detect **a)** GFP **b)** c-Myc tag for Her2 CAR and **c)** Mcl-1. β-actin was used as control for the Western blots. All representative blots above are repeated three times and quantified and presented in the bar graph (right) using Image Studio Lite. Bar graph values represent the mean values ± SD from three different repeats.

**Fig. 3.**
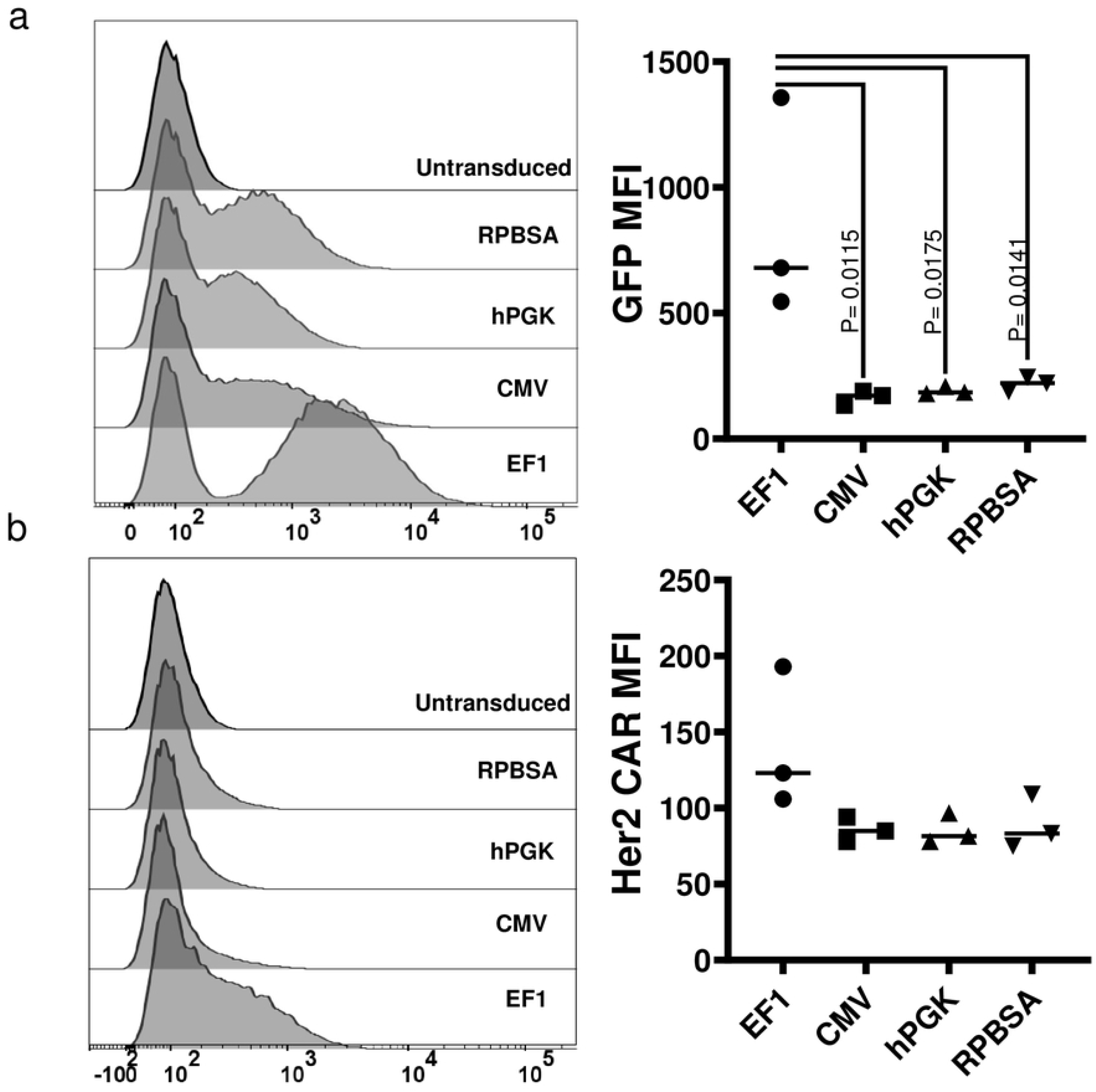
GFP and Her2 CAR expression by four constructs in primary T cells. Flow cytometry carried out to measure the expression of **a)** GFP and **b)** Her2 CAR stained for c-myc tag. Dead cells were excluded by Zombie NIR viability dye at analysis. MFI for three different donors are shown in graphs.

### Functional effect of CAR T cells in tumour and T cell engagement

We next examined the function of the CAR T cells transduced with each of the promoter constructs, measuring cytokine release (IL-2 and IFN-γ), cytotoxicity and activation following incubation of CAR T cells with the Her2+ MCF-7 breast cancer cell line. Although the expression of CD69 as an activation marker was similarly expressed among the CAR T cells with different promoters (Fig. 4a), EF-1 and CMV CAR T cells showed optimal cytokine release after engaging MCF-7 cells (Fig. 4b & c). CAR T cells transduced with hPGK were less active and those with the RPBSA construct failed to release detectable IL-2 and IFN-γ. Cytotoxicity assay with the four constructs showed similar results with strong killing with CAR T cells expressing under the EF-1 promoter at 24 h time points (Fig. 4d).

**Fig. 4.**
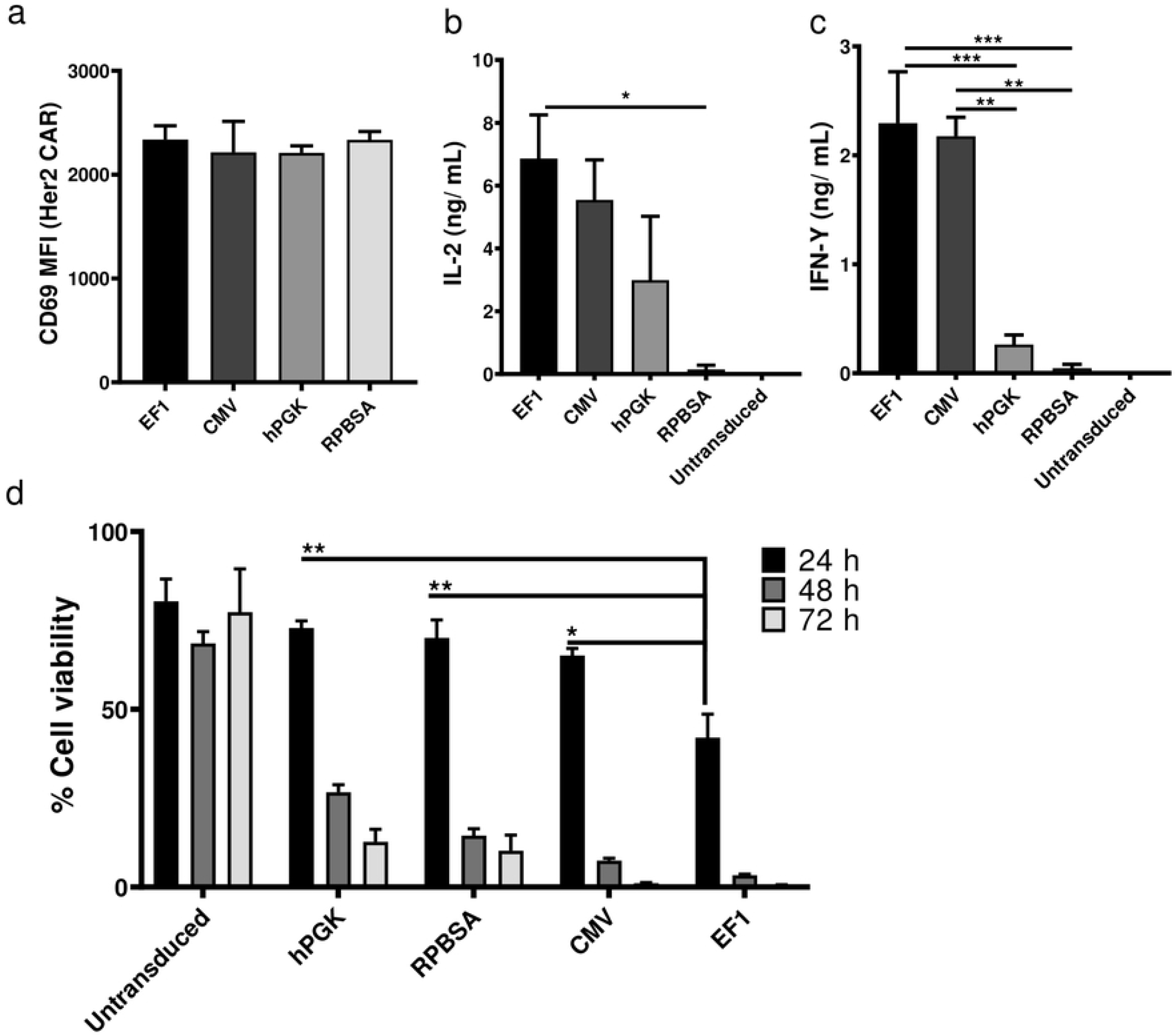
Comparison of anti-tumor activity between promoters in CAR T cells. **a)** Flow cytometric analysis of Her2 CAR T cells 18 h after co-culture with Her2+ / MCF-7 cells. Data shows the MFI of CD69 expression from three different donors. Bar graphs show the secretion of **b)** IL-2 and **c)** IFN-γ by different CAR T cells measured by ELISA. CAR T cells were incubated with Her-2+ MCF-7 cell line for 24 h before supernatants were collected. Cytokines were measured in ng/mL. **d)** Luciferase based cytotoxicity assay assessed 24, 48 and after 72 h after incubation of CAR T cells with MCF-7 cells stably expressing the firefly luciferase gene. The graph shows the percentage of cell viability, calculated by dividing the luciferase of the sample well over the luciferase reading of untreated MCF-7.

To functionally test the relationship between the expression level of the most distal gene Mcl-1, and resistance to AICD, CAR T cells carrying four different promoters were challenged with 1 μg/mL of LZ-CD95L and mitochondrial depolarisation monitored by TMRE staining and flow cytometry. Again, EF-1 provided the most potent protection against CD95L-induced cell death (Fig. 5).

**Fig. 5.**
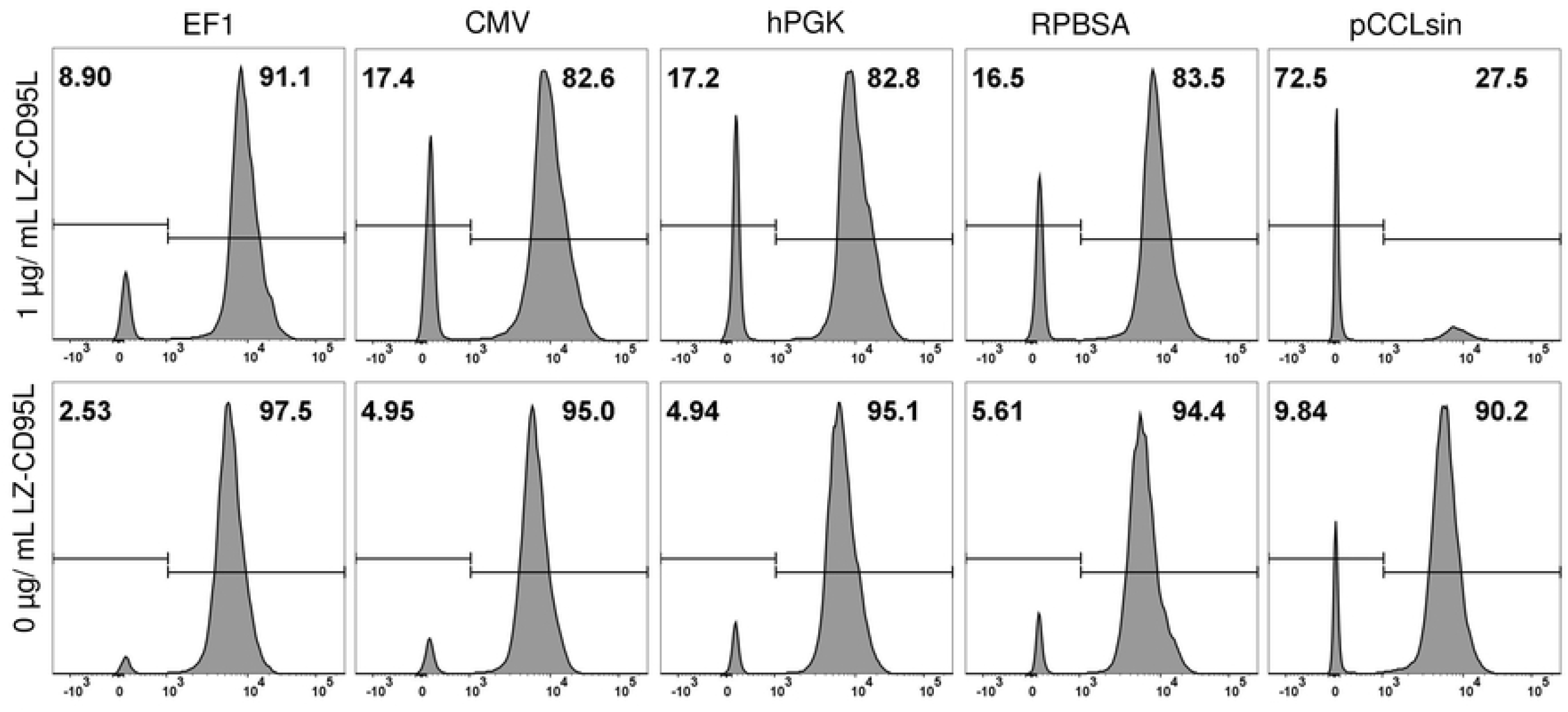
TMRE assay for monitoring mitochondrial membrane potential. CAR T cells expressing Mcl-1 as an anti-apoptotic gene as the most distal gene in the cassette were challenged with 1 μg/mL (top) or 0 μg/mL (below) of LZ-CD95L to mimic AICD. The right peak in the histograms show the intact healthy mitochondria and left peak are depolarised mitochondria. pCCLsin (lentivector expressing only GFP) was used as control.

### Promoter comparison for driving short transcripts

We compared the ability of the four promoters in transcribing GFP linked to an FMC63 CD19 CAR, the most studied CAR construct and the first CAR T cell design approved by the FDA. The FMC63 CAR transcript is 1.2 kb shorter than the GFP-Her2 CAR-Mcl1. Viral titres and transduction efficacies were similar among all promoters driving the shorter FMC63 CAR mRNA (Fig. 6a & b). Protein expression of the shorter GFP-CAR constructs was superior in HEK293T transduced with EF-1 and CMV constructs (Fig. 6c). In primary T cells, EF-1 gave the highest expression for GFP and CD19 CAR, while CMV gave a more heterogenous expression, but this was not statistically significantly lower than EF-1 (Fig. 6d & e).

**Fig. 6.**
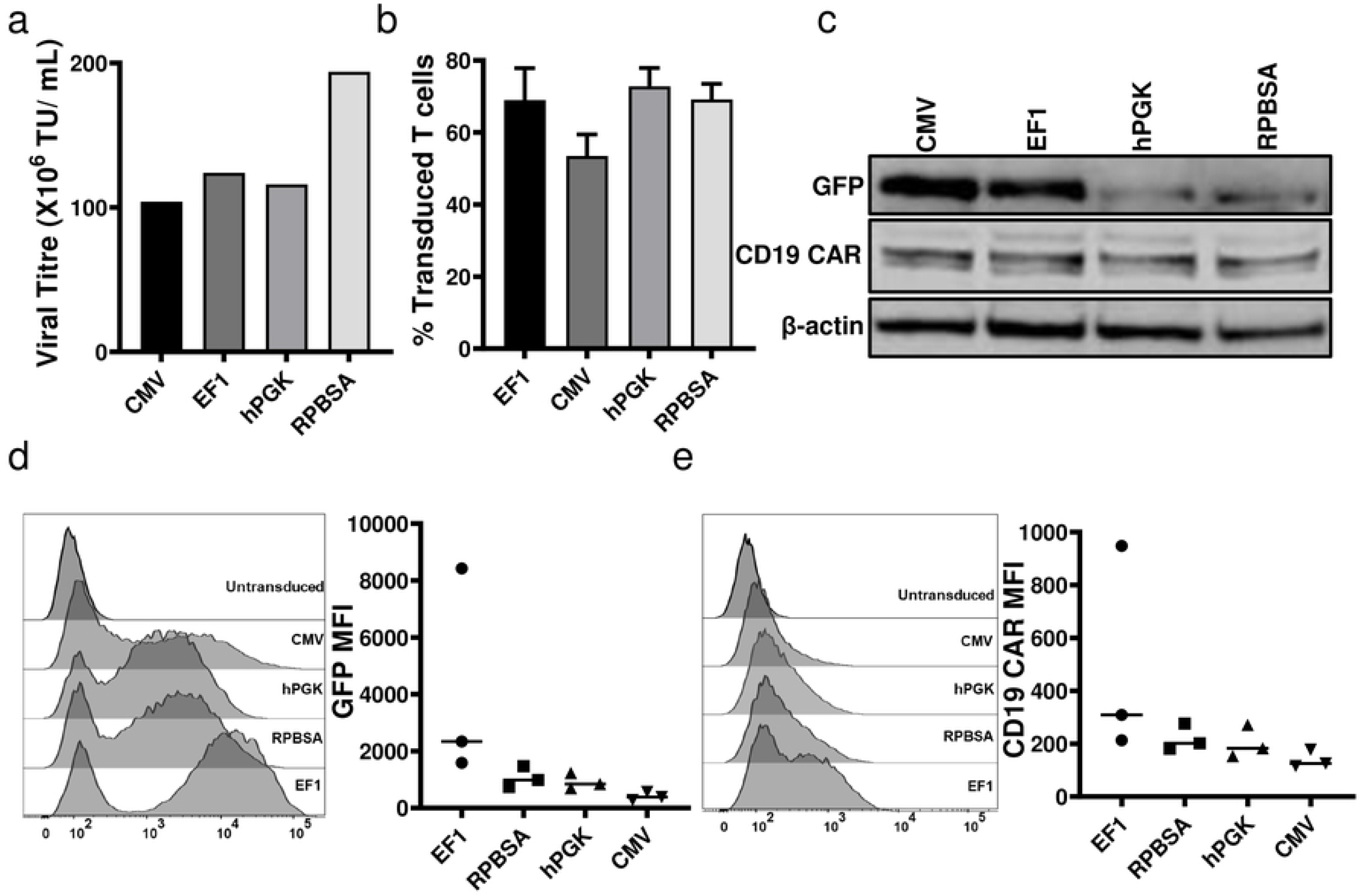
Comparison of four constructs for transcribing short RNA. The eGFP gene linked to FMC63 CD19 CAR was cloned under the control of the four promoters. **a)** Titration and **b)** transduction efficacy among four constructs **c)** Western blot analysis for GFP and CD19 CAR level of HEK293T cells transduced at MOI 2:1 plus 1 μg/mL polybrene **d)** GFP and **e)** CD19 CAR expression in CAR T cells by flow cytometry.

Although CD69 expression on antigen stimulated CAR T cells was similar for all promoter constructs (Fig. 7a), EF-1 constructs drove higher levels of CAR triggering in terms of cytokine release and cytotoxicity. Interestingly, RPBSA showed optimal function in CAR T cells expressing the shorter CD19 CAR transcript (Fig. 7 b, c & d) emphasizing that promoter activity is dependent on the nature of the downstream transcript.

**Fig. 7.**
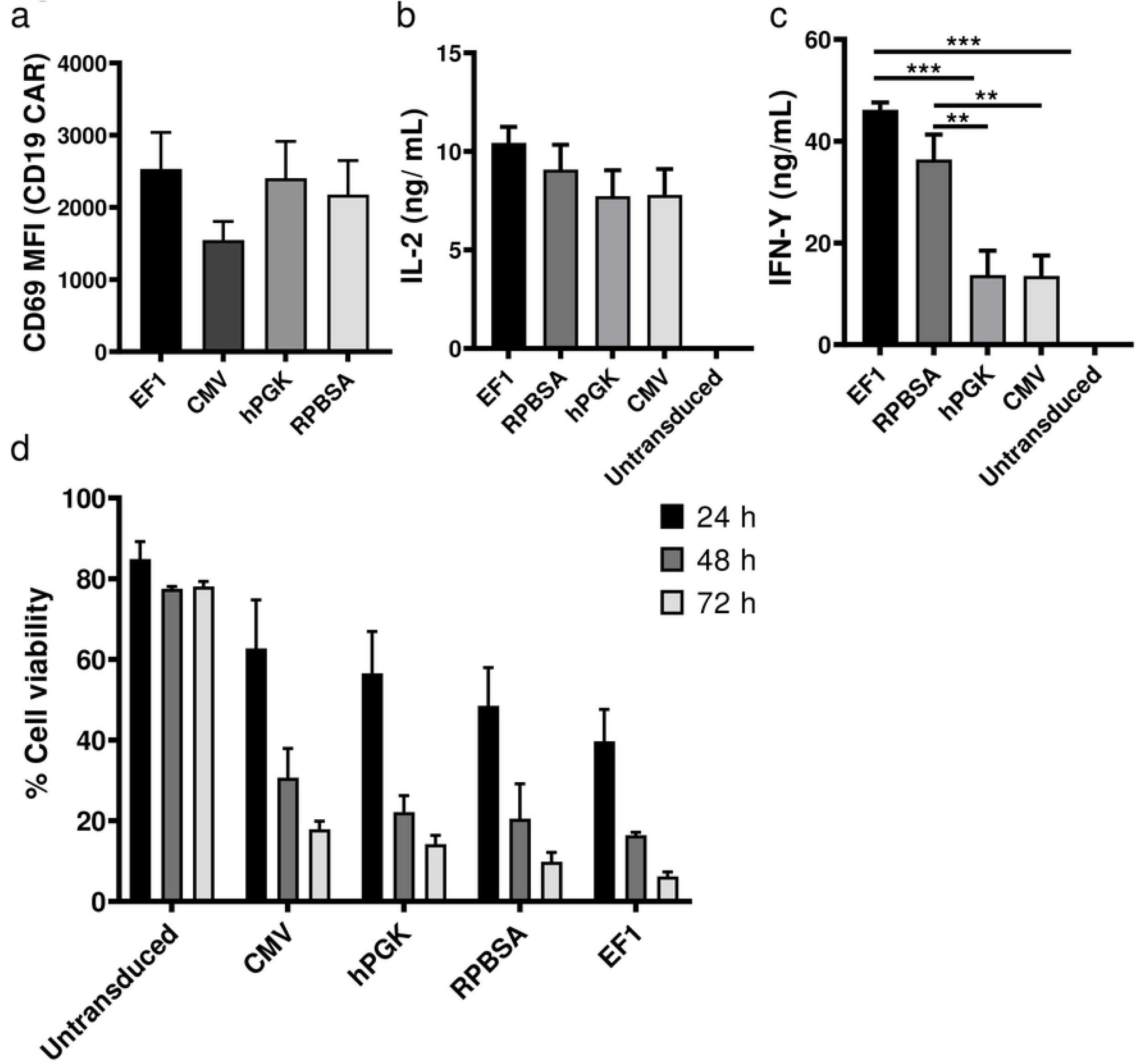
Functional analysis of CAR T cells expressing short RNA. **a)** CD69 activation assay was carried out 18 hours after incubation of the four types of promoter-driven CD19 CAR T cells with CD19+ HEK293T cells. **b & c)** Cytokine release assay for secretion of **b)** IL-2 and **c)** IFN-γ. CAR T cells were co-cultured with CD19+ HEK239T and supernatant were collected after 24 h. **d)** Luciferase based cytotoxicity assay assessed 24, 48 and after 72 h after incubation of CD19 CAR T cells with CD19+ HEK293 cells stably expressing the firefly luciferase gene. The graph shows the percentage of cell viability, calculated by dividing the luciferase of sample well over the luciferase reading of untreated HEK293T.

### Core promoter elements, CpG island and TF binding sites are varying between promoters

Although all four selected promoters are assumed to be constitutive and active in most cell types, bioinformatic analysis showed that the four promoters vary in terms of core promoter elements and potential TF binding sites. While there is no universal core promoter elements for RNA polymerase II, the TATA box, initiator (Inr) element, TFIIB recognition element (BRE), downstream core promoter element (DPE) and motif ten element (MTE) are well-established core promoter elements (Fig. 8a). Overall, EF-1 had more core promoter elements, such as GC box, DPE and MTE (Fig. 8b, Table 2). Except for hPGK, all promoters contain a TATA box.

**Fig. 8.**
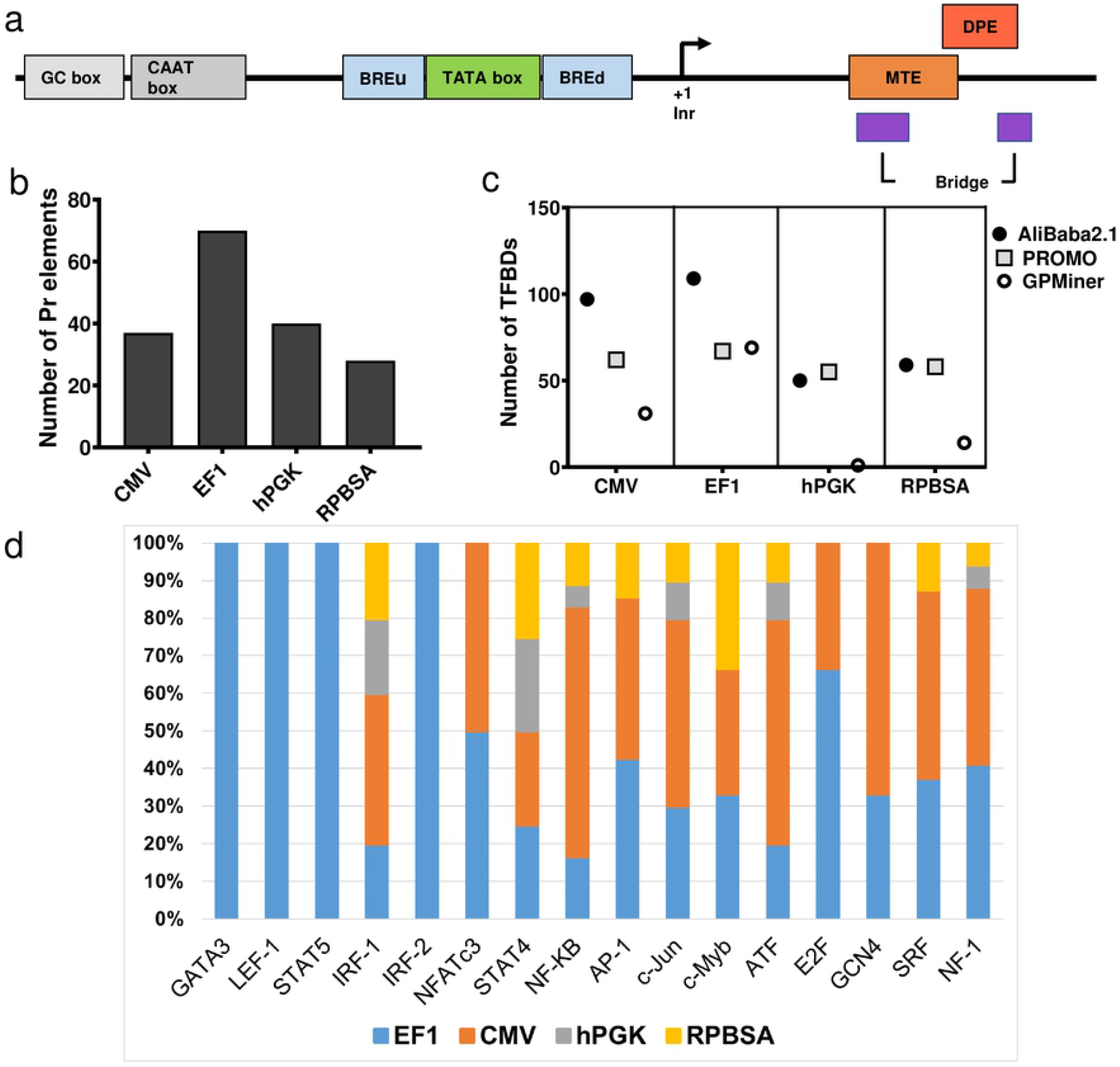
Structure and bioinformatics analysis of four different promoters **a)** Structure of a eukaryotic core promoter and the position of core elements within a promoter were investigated in four different promoters **b)** Total number of core promoter elements predicted by YAPP, GPMiner and ElemeNT algorithms (details provided in Table 2) **c)** The number of TF binding sites in promoters sequenced analyzed by AliBaba2.1, PROMO and GPMiner programs d) Enrichment of sixteen TFs in four promoters highly expressed in T cells. The data shows the percentage of total numbers predicted for binding sites.

Another feature of eukaryotic promoters is the presence of CpG islands. CpG islands could result in hypermethylation and gene silencing. However, promoters with CpG islands containing multiple Sp1 binding sites exhibit a hypomethylated state and are typically stronger promoters (25). We therefore searched for CpG islands within our promoters using two different programs (Table 1). Except for CMV, all promoters were expected to have at least one CpG island. When we searched the Sp1 binding sites within the CpG islands, EF-1 and hPGK showed the highest number of Sp1 binding sites in their CpG islands (Table 3). EMBOSS Cpgplot program predicted two CpG island for EF-1 with 37 Sp1 binding sites. Lastly, looking at individual TFs and their expression in T cells, sixteen TFs were selected based on their function and expression in T cells (26–28). EF-1 showed to have binding sites for all these TFs. CMV is the next promoter enriched for these promoter-specific TFs excluding GATA3, LEF-1, STAT5 and IF-2 (Fig. 8d).

## Discussion

In this study, we compared four promoters for optimal expression of long RNA encoding multiple gene products in CAR T cells. Our results suggest that promoter requirements are stringent for driving long RNA, and that EF-1 is the best choice for driving short or long RNA in CAR T cells, similar to an early study (29). In contrast to the poor results obtained here for hPGK and RPBSA in driving long and complex RNA, these same promoters demonstrate little difference to the so-called strong promoters CMV and EF-1 lentiviral based systems to drive shorter RNA sequences, such as CAR and fluorescent reporter genes or a simpler GFP (see Fig. 6e) – consistent with other studies (1, 8, 9, 21, 23).

To determine the functional role of additional accessory genes expressed in long constructs, we utilised Mcl-1, a bcl2 family member with an essential role in T cell development, mitochondrial function and lifespan. To our knowledge, this is the first study to demonstrate that Mcl-1 is a suitable candidate for enhancing CAR T cell performance (15, 16). Expression of mcl1 in a position distal to the CAR allowed protection from CD95-induced cell death. Interestingly, although protection was noted with all promoters, EF-1 driven-cassettes consistently gave the best protection. The fact that protection was observed with mcl1 driven by the weaker promoters RPBSA and hPGK contrasts with the stringent requirement for a strong promoter to drive CAR expression for optimal cytotoxicity and cytokine release.

Our analysis of promoter motifs demonstrates clear differences in transcription factor binding sites and core promoter elements between the strong (EF-1 and CMV) and weaker (hPGK and RPBSA) promoters. Although not all the predicted core promoter elements might be functional in primary T cells, the high number of the core elements can correlate with the strength of the promoter (25). In addition EF-1 and CMV predominantly enriched for TFs specific or highly expressed in T cells (26–28, 30, 31) such as GATA3, NFATc3, NF-kB, AP1 and c-Jun (Fig. 1e), The number of transcription factor and core promoter element sites predicted within the promoters may provide some explanation for the ability of the CMV and EF-1 promoters to direct long mRNA expression (Fig. 1, Supplementary data).

The activity of promoters with predicted ‘ubiquitous’ expression, such as the four studied here, will still depend greatly on the lineage of the host cell (32). However, EF-1 promoter was found to be active and resistant to silencing in cells where other viral promoters may become silenced (33). Therefore, future work will be required to determine if the superior performance of EF-1 and CMV in expressing long RNA sequences can be extrapolated to other cell primary cell types.

In our study, the lower expression of CAR within a long mRNA transcript driven by the RPBSA and hPGK translated into lower lytic function for a Her2-expressing tumor cell line. Given the profound effects that CAR density has on T cell activation, our results will be useful for developing strategies to titrate CAR expression at the T cell. Promoter choice would be expected to be a critical consideration for controlling the levels of surface expressed CAR, which in turn would dictate the level of T cell activation, lytic function, as well as undesirable tonic (antigen-independent) signaling (2, 34–37). Optimal CAR expression will be critical for minimizing tonic signaling, while optimizing signal transduction during antigen-specific signaling. In addition, lowering the level of CAR expression could contribute desirable avidity effects to T cell recognition of antigen, thereby minimizing CAR T cell activation by tumor-associated antigen on self-tissue (14). Interestingly, despite CMV inducing a noticeably higher level expression of GFP, CAR and Mcl-1 in HEK293T cells, as compared to EF-1, functional analysis showed superior activation of primary human CAR T cells driven by EF-1 in terms of cytokine release and cytotoxicity against MCF-7. In addition EF-1 driven expression of Mcl-1 provided the best protection of CAR T cells to AICD induced by CD95L.

A further consideration for promoter choice is possible silencing *in vivo.* In particular, CMV can be silenced after a period of weeks post-transduction (32, 38). However, the effects of promoter silencing might be overshadowed by the long term CAR T cell downregulation that occurs in a methylation-independent fashion following CAR triggering both *in vitro* and *in vivo* (14, 39, 40). In conclusion, the study of long mRNA production will improve our ability to express multiple genes in CAR T cells to improve cell survival and persistence of infused CAR T cells.

## Conflict of interest

The authors declare that they have no conflicts of interest with the contents of this article.

## Author Contribution

AR and AM wrote the paper and supervised the study. AR performed or contributed to experimental data in all figures, prepared the tables and performed the analysis of the promoter regions. GT and AP assisted with planning, experiments and contributed to the writing of the paper.

## Acknowledgments

We thank Dr. Sarah Saunderson for critical review of the manuscript.

## References

1. Milone MC, Fish JD, Carpenito C, Carroll RG, Binder GK, Teachey D, et al. Chimeric receptors containing CD137 signal transduction domains mediate enhanced survival of T cells and increased antileukemic efficacy in vivo. Molecular therapy: the journal of the American Society of Gene Therapy. 2009;17(8):1453–64.

2. Frigault MJ, Lee J, Basil MC, Carpenito C, Motohashi S, Scholler J, et al. Identification of chimeric antigen receptors that mediate constitutive or inducible proliferation of T cells. Cancer immunology research. 2015;3(4):356–67.

3. Drent E, Poels R, Mulders MJ, van de Donk N, Themeli M, Lokhorst HM, et al. Feasibility of controlling CD38-CAR T cell activity with a Tet-on inducible CAR design. PloS one. 2018;13(5):e0197349.

4. Stoiber S, Cadilha BL, Benmebarek MR, Lesch S, Endres S, Kobold S. Limitations in the Design of Chimeric Antigen Receptors for Cancer Therapy. Cells. 2019;8(5).

5. Naldini L, Trono D, Verma IM. Lentiviral vectors, two decades later. Science (New York, NY). 2016;353(6304):1101–2.

6. McLellan AD, Ali Hosseini Rad SM. Chimeric antigen receptor T cell persistence and memory cell formation. Immunology and cell biology. 2019.

7. Gilham DE, Lie ALM, Taylor N, Hawkins RE. Cytokine stimulation and the choice of promoter are critical factors for the efficient transduction of mouse T cells with HIV-1 vectors. The journal of gene medicine. 2010;12(2):129–36.

8. Jones S, Peng PD, Yang S, Hsu C, Cohen CJ, Zhao Y, et al. Lentiviral vector design for optimal T cell receptor gene expression in the transduction of peripheral blood lymphocytes and tumor-infiltrating lymphocytes. Human gene therapy. 2009;20(6):630–40.

9. Amendola M, Venneri MA, Biffi A, Vigna E, Naldini L. Coordinate dual-gene transgenesis by lentiviral vectors carrying synthetic bidirectional promoters. Nature biotechnology. 2005;23(1):108–16.

10. Curtin JA, Dane AP, Swanson A, Alexander IE, Ginn SL. Bidirectional promoter interference between two widely used internal heterologous promoters in a late-generation lentiviral construct. Gene therapy. 2008;15(5):384–90.

11. Wang D, Shao Y, Zhang X, Lu G, Liu B. IL-23 and PSMA-targeted duo-CAR T cells in Prostate Cancer Eradication in a preclinical model. Journal of translational medicine. 2020;18(1):23.

12. Hoyos V, Savoldo B, Quintarelli C, Mahendravada A, Zhang M, Vera J, et al. Engineering CD19-specific T lymphocytes with interleukin-15 and a suicide gene to enhance their anti-lymphoma/leukemia effects and safety. Leukemia. 2010;24(6):1160–70.

13. Philip B, Kokalaki E, Mekkaoui L, Thomas S, Straathof K, Flutter B, et al. A highly compact epitope-based marker/suicide gene for easier and safer T-cell therapy. Blood. 2014;124(8):1277–87.

14. Eyquem J, Mansilla-Soto J, Giavridis T, van der Stegen SJ, Hamieh M, Cunanan KM, et al. Targeting a CAR to the TRAC locus with CRISPR/Cas9 enhances tumour rejection. Nature. 2017;543(7643):113–7.

15. Andersen JL, Kornbluth S. Mcl-1 rescues a glitch in the matrix. Nat Cell Biol. 2012;14(6):563–5.

16. Morciano G, Pedriali G, Sbano L, Iannitti T, Giorgi C, Pinton P. Intersection of mitochondrial fission and fusion machinery with apoptotic pathways: Role of Mcl-1. Biol Cell. 2016;108(10):279–93.

17. Kim EH, Neldner B, Gui J, Craig RW, Suresh M. Mcl-1 regulates effector and memory CD8 T-cell differentiation during acute viral infection. Virology. 2016;490:75–82.

18. Dzhagalov I, Dunkle A, He YW. The anti-apoptotic Bcl-2 family member Mcl-1 promotes T lymphocyte survival at multiple stages. Journal of immunology (Baltimore, Md: 1950). 2008;181(1):521–8.

19. Dunkle A, Dzhagalov I, He YW. Cytokine-dependent and cytokine-independent roles for Mcl-1: genetic evidence for multiple mechanisms by which Mcl-1 promotes survival in primary T lymphocytes. Cell death & disease. 2011;2:e214.

20. Stewart DP, Koss B, Bathina M, Perciavalle RM, Bisanz K, Opferman JT. Ubiquitin-independent degradation of antiapoptotic MCL-1. Mol Cell Biol. 2010;30(12):3099–110.

21. Dull T, Zufferey R, Kelly M, Mandel RJ, Nguyen M, Trono D, et al. A third-generation lentivirus vector with a conditional packaging system. Journal of virology. 1998;72(11):8463–71.

22. Fu X, Tao L, Rivera A, Williamson S, Song X-T, Ahmed N, et al. A simple and sensitive method for measuring tumor-specific T cell cytotoxicity. PloS one. 2010;5(7).

23. Kowarz E, Loscher D, Marschalek R. Optimized Sleeping Beauty transposons rapidly generate stable transgenic cell lines. Biotechnol J. 2015;10(4):647–53.

24. Yee J-K, Moores JC, Jolly DJ, Wolff JA, Respess JG, Friedmann T. Gene expression from transcriptionally disabled retroviral vectors. Proceedings of the National Academy of Sciences. 1987;84(15):5197–201.

25. Butler JE, Kadonaga JT. The RNA polymerase II core promoter: a key component in the regulation of gene expression. Genes & development. 2002;16(20):2583–92.

26. Naito T, Tanaka H, Naoe Y, Taniuchi I. Transcriptional control of T-cell development. Int Immunol. 2011;23(11):661–8.

27. Hosokawa H, Rothenberg EV. Cytokines, Transcription Factors, and the Initiation of T-Cell Development. Cold Spring Harb Perspect Biol. 2018;10(5).

28. Rothenberg EV. The chromatin landscape and transcription factors in T cell programming. Trends Immunol. 2014;35(5):195–204.

29. Frigault MJ, Lee J, Basil MC, Carpenito C, Motohashi S, Scholler J, et al. Identification of chimeric antigen receptors that mediate constitutive or inducible proliferation of T cells. Cancer immunology research. 2015;3(4):356–67.

30. Urso K, Alfranca A, Martinez-Martinez S, Escolano A, Ortega I, Rodriguez A, et al. NFATc3 regulates the transcription of genes involved in T-cell activation and angiogenesis. Blood. 2011;118(3):795–803.

31. Lynn RC, Weber EW, Sotillo E, Gennert D, Xu P, Good Z, et al. c-Jun overexpression in CAR T cells induces exhaustion resistance. Nature. 2019;576(7786):293–300.

32. Hong S, Hwang DY, Yoon S, Isacson O, Ramezani A, Hawley RG, et al. Functional analysis of various promoters in lentiviral vectors at different stages of in vitro differentiation of mouse embryonic stem cells. Molecular therapy: the journal of the American Society of Gene Therapy. 2007;15(9):1630–9.

33. Wang X, Xu Z, Tian Z, Zhang X, Xu D, Li Q, et al. The EF-1alpha promoter maintains high-level transgene expression from episomal vectors in transfected CHO-K1 cells. J Cell Mol Med. 2017;21(11):3044–54.

34. Chang ZL, Silver PA, Chen YY. Identification and selective expansion of functionally superior T cells expressing chimeric antigen receptors. Journal of translational medicine. 2015;13:161.

35. Chmielewski M, Hombach A, Heuser C, Adams GP, Abken H. T cell activation by antibody-like immunoreceptors: increase in affinity of the single-chain fragment domain above threshold does not increase T cell activation against antigen-positive target cells but decreases selectivity. Journal of immunology (Baltimore, Md: 1950). 2004;173(12):7647–53.

36. Han C, Sim SJ, Kim SH, Singh R, Hwang S, Kim YI, et al. Desensitized chimeric antigen receptor T cells selectively recognize target cells with enhanced antigen expression. Nature communications. 2018;9(1):468.

37. Sakemura R, Terakura S, Watanabe K, Julamanee J, Takagi E, Miyao K, et al. A Tet-On Inducible System for Controlling CD19-Chimeric Antigen Receptor Expression upon Drug Administration. Cancer immunology research. 2016;4(8):658–68.

38. Brooks AR, Harkins RN, Wang P, Qian HS, Liu P, Rubanyi GM. Transcriptional silencing is associated with extensive methylation of the CMV promoter following adenoviral gene delivery to muscle. The journal of gene medicine. 2004;6(4):395–404.

39. Burns WR, Zheng Z, Rosenberg SA, Morgan RA. Lack of specific gamma-retroviral vector long terminal repeat promoter silencing in patients receiving genetically engineered lymphocytes and activation upon lymphocyte restimulation. Blood. 2009;114(14):2888–99.

40. Gallegos AM, Xiong H, Leiner IM, Susac B, Glickman MS, Pamer EG, et al. Control of T cell antigen reactivity via programmed TCR downregulation. Nature immunology. 2016;17(4):379–86.

